# Prefrontal Transcranial Direct Current Stimulation globally improves learning, but does not selectively potentiate the benefits of Targeted Memory Reactivation on awake memory consolidation

**DOI:** 10.1101/2020.04.13.039214

**Authors:** Médhi Gilson, Michael A. Nitsche, Philippe Peigneux

## Abstract

Targeted Memory Reactivation (TMR) and transcranial Direct Current Stimulation (tDCS) can enhance memory consolidation. It is currently unknown whether TMR reinforced by simultaneous tDCS has superior efficacy. In this study, we investigated the complementary effect of TMR and bilateral tDCS on the consolidation of emotionally neutral and negative declarative memories. Participants learned neutral and negative word pairs. Each word pair was presented with an emotionally compatible sound. Following learning, participants spent a 20-minutes retention interval awake under 4 possible conditions: (1) TMR alone (i.e. replay of 50% of the associated sounds), (2) TMR combined with anodal stimulation of the left DLPFC, (3) TMR combined with anodal stimulation of the right DLPFC and (4) TMR with sham tDCS. Results evidenced selective memory enhancement for the replayed stimuli in the TMR-only and TMR-sham conditions, which confirms a specific effect of TMR on memory. However, memory was enhanced at higher levels for all learned items (irrespective of TMR) in the TMR-anodal right and TMR-anodal left tDCS conditions, suggesting that the beneficial effects of tDCS overshadow the specific effects of TMR. Emotionally negative memories were not modulated by tDCS hemispheric polarity. We conclude that electrical stimulation of the DLPFC during post-learning period globally benefits memory consolidation, but does not potentiate the specific benefits of TMR.

## 1. Introduction

Memory consolidation is the process by which novel and fragile memory traces are progressively transformed into more robust representations (McGaugh, 1966). Consolidation is possibly supported by the offline replay of the neuronal activity that subtends learning processes (Buzsaki, 1996). Supporting this hypothesis, animal and human studies evidenced the reinstatement of learning-related brain activity during offline post-training periods, both at wake (Humiston, Tucker, Summer, & Wamsley, 2019; Peigneux et al., 2006; Tambini & Davachi, 2019; Wamsley, 2019) and during sleep (Jegou et al., 2019; Peigneux et al., 2004; Sara, 2010; Schonauer et al., 2017; Skaggs & Mcnaughton, 1996; Valdes, McNaughton, & Fellous, 2015). Although several studies reported better consolidation in declarative memory after sleep than wakefulness, mostly associated with the triggering of learning-related brain activity during slow wave sleep (SWS; for reviews see e.g. Born & Wilhelm, 2012; S. Diekelmann & Born, 2010; Hu, Cheng, Chiu, & Paller, 2020; Westermann, Lange, Textor, & Born, 2015), others showed that quiet wakefulness can also benefit memory consolidation (e.g., Craig, Ottaway, & Dewar, 2018; Dewar, Alber, Butler, Cowan, & Della Sala, 2012; Dewar, Garcia, Cowan, & Della Sala, 2009; Humiston et al., 2019; Tambini & Davachi, 2019; Tambini, Ketz, & Davachi, 2010; Wamsley, 2019). For instance, memory for stories was found superior after 10 minutes spent in a wakeful resting state than after an equivalent period of time spent in active wakefulness (Dewar et al., 2012). To some extent, the neural oscillatory mechanisms involved in memory formation at wake might share similarities with sleep (Buzsaki, 1989). Accordingly, Tambini and colleagues (Tambini et al., 2010) found that enhanced hippocampal-cortical coordination during wakeful rest after learning is predictive of memory performance. Furthermore, increased slow oscillatory activity (1 Hz) and reduced alpha (8-12 Hz) activity during quiet resting were associated with memory improvement for learned stories, suggesting that slow oscillations at wake might promote the dialogue between hippocampal and cortical regions (Brokaw et al., 2016), like during sleep (Buzsaki, 1996). Therefore, post-training resting wakefulness might improve memory performance not only by reducing ongoing interferences, but also by providing a neural milieu that favours the interactions between subcortical hippocampal areas and cortical regions subtending memory consolidation, including prefrontal areas (Wamsley, 2019).

Targeted Memory Reactivation (TMR), i.e. the presentation of learning-related cues during offline periods, can additionally enhance memory consolidation (for reviews see e.g. Hu et al., 2020; Oudiette & Paller, 2013). For instance, TMR during post-training wakefulness was found to rescue targeted memories from forgetting (Farthouat, Gilson, & Peigneux, 2017; Oudiette, Antony, Creery, & Paller, 2013), although this was not always replicated (see e.g. Schreiner & Rasch, 2015b and Wilhelm, Schreiner, Beck, & Rasch, 2020 for a lack of effect on vocabulary learning). TMR was also found detrimental in putting memories in a labile state and increasing sensitivity to interference (Diekelmann, Buchel, Born, & Rasch, 2011), and in some cases was more (Oudiette et al., 2013) or solely (Rudoy, Voss, Westerberg, & Paller, 2009; Schreiner & Rasch, 2015a) beneficial during sleep (for a review see e.g. Lewis & Bendor, 2019). Besides, another promising technique to promote learning and memory consolidation processes is transcranial Direct Current Stimulation (tDCS) that modulates cortical excitability (Brunoni et al., 2012; Huang et al., 2017; Kuo & Nitsche, 2012; Nitsche & Paulus, 2000). Anodal excitatory stimulation of the left dorsolateral prefrontal cortex (DLPFC) during encoding improves declarative memory (Javadi & Walsh, 2012; Westphal et al., 2019), whereas cathodal inhibitory stimulation exerts a detrimental effect (Javadi & Walsh, 2012; Lang, Nitsche, Paulus, Rothwell, & Lemon, 2004). Similarly, cathodal stimulation of the left DLPFC impairs recognition performance, whereas it tends to be improved by anodal stimulation. Furthermore, tDCS may also benefit offline consolidation mechanisms. Indeed, although 20 minutes of anodal stimulation over the temporoparietal cortex did not modify the rate of learning in elderly participants, it enhanced delayed free recall performance as measured one week later (Floel et al., 2012). Similarly, anodal stimulation over the premotor cortex during rapid eye movement sleep (Nitsche et al., 2010) or at wake after learning (Krause, Meier, Dinkelbach, & Pollok, 2016) was shown to benefit the consolidation of a motor sequence

Finally, emotional and arousing memories are usually better remembered than memories without any affective load (for a review see (Deliens, Gilson, & Peigneux, 2014). A potential hemispheric lateralization in the processing of emotion has been proposed. According to the right hemisphere hypothesis, the right hemisphere would be involved in the general processing of emotions (Borod, Bloom, Brickman, Nakhutina, & Curko, 2002). In contrast, the valence specific hypothesis posits that the left hemisphere is dominant for positive emotions and the right hemisphere for negative ones (Adolphs, Jansari, & Tranel, 2001; Balconi & Ferrari, 2013).

To the best of our knowledge, it is currently unknown whether TMR reinforced by simultaneous tDCS during the offline wake period has superior efficacy for the consolidation of verbal declarative memories. In the present study, we tested this hypothesis using TMR alone or in combination with tDCS over the dorsolateral prefrontal cortex (DLPFC) after the learning of pairs of words. Additionally, we investigated the effect of tDCS lateralization on the consolidation of emotionally neutral and negative word pairs, considering a potential hemispheric lateralization in the processing of emotions (Adolphs et al., 2001; Borod et al., 1998). Based on the right hemisphere and the valence specific hypotheses, we expected the combination of excitatory anodal stimulation over the right DLPFC and inhibitory cathodal stimulation over the left DLPFC to benefit more the consolidation of negative memories than tDCS with reversed polarity (left anodal/right cathodal).

In the present study, healthy young participants learned a list of neutral and negative word pairs. Each word pair was associated with an emotionally compatible sound at learning. Participants then spent 20 minutes awake in a quiet environment in one out of four possible stimulation conditions, i.e. TMR-only (half of the sounds associated with the word pairs to remember replayed during the 20-minute consolidation interval), TMR-anodal left tDCS (TMR plus anodal stimulation on left DLPFC and cathodal stimulation on right DLPFC), TMR-anodal right tDCS (identical with reversed polarity), and TMR-sham tDCS (TMR plus tDCS for 15 seconds only at the beginning of the 20-minute period). We predicted better memory performance for cued word pairs than for non-cued word pairs, and that this effect would be potentiated by tDCS. Additionally, we expected a specific modulation of negative memories in the TMR-anodal right tDCS condition.

## 2. Material and methods

### 2.1. Participants

Seventy-two healthy participants gave their written informed consent to participate in this study approved by the Faculty Ethics committee at the Université Libre de Bruxelles (ULB). Three of them were excluded because they did not participate in the entirety of the experiment. The sixty-nine remaining participants were native French speakers, right-handed, free of medication known to influence sleep quality and/or mood. They reported not suffering or having suffered from neurological or psychiatric disorders. Pre-study examination evidenced alexithymia scores below cutoff score (mean ± standard deviation 41.08 ± 9.2, cut-off score 61; Alexithymia Toronto Scale, Bagby, Parker, & Taylor, 1994) and vocabulary knowledge within normative values (score range 25–40/44; Mill-Hill Vocabulary Scale, Deltour, 1993). Participants were randomly assigned to one out of four conditions.

### 2.2. Material

A neutral and a negative list of unrelated French word pairs (18 pairs/list) were selected based on their emotional valence (neutral = 16.19 ± 13.7 vs. negative = 77.67 ± 19.8, *t*(1, 70) = 15.33, *p* < .0001; Syssau & Font, 2005). The two lists were equated according to lexical frequency (neutral = 54.06 ± 67.5 vs. negative = 56.14 ± 86.2, *t*(1, 70) = 0.11, *p*= .91; Content, Mousty, & Radeau, 1990), imaging valence (neutral = 5.43 ± 1.6 vs. negative = 4.85 ± 1.5, *t*(1, 70) = −1.55, *p* = .12; Desrochers & Bergeron, 2000), number of syllables (neutral = 1.75 ± 0.7 vs. negative = 1.97 ± 0.7, *t*(1, 70) = 1.32, *p* = .19) and number of letters (neutral = 5.86 ± 1.5 vs. negative = 6.38 ± 1.8, *t*(1, 70) = 1.34, *p* = .19).

Each word pair was randomly associated with a specific sound (duration 6 sec), taken from the International Affective Digitized Sounds database (IADS, Bradley and Lang, 2007). Neutral word pairs were matched with neutral sounds and negative word pairs were matched with negative sounds. Negative sounds were selected based on high arousal and low pleasantness ratings on a 9-point Likert scale (Bradley & Lang, 1994), while neutral sounds were selected according to middle arousal and middle pleasantness ratings (mean IADS pleasure rating neutral = 4.94 ± 0.21 vs. negative = 2.12 ± 0.38, *t*(1, 34) = 26.92, *p* < .001; mean IADS arousal rating neutral = 4.93 ± 0.8 vs. negative = 7.34 ± 0.5, *t*(1, 34) = −10.48, *p* < .001).

### 2.3. Procedure

The experimental procedure is illustrated in Figure 1. All participants started with the Encoding phase during which each of the 36 word pairs was displayed one by one on a computer screen for 6 seconds while the associated sound was delivered through headphones. Between each word pair, a yellow fixation cross was displayed for 4 seconds. It turned red 1 second before the apparition of the next pair. The 36 word pairs were presented twice in random order. Participants were then administered an Immediate cued Recall Test (IRT): the first word of each pair was displayed on the screen together with its associated sound, and participants had to type in the associated word. The correct response was then displayed on screen to promote error-free learning. If the participant’s total recall score was below the minimum criterion of 75% of correct responses, the incorrectly recalled word pairs were represented and then the IRT administered again on all pairs, until participants’ recall performance was at least 75%. After this learning session, participants were seated in a comfortable chair and asked to rest quietly for 20 minutes (wakeful resting period) while listening to series of sounds. They were pseudo-randomly assigned to one out of four possible conditions. In the TMR-only condition (N = 16 participants), half of the sounds (9 neutral and 9 negative) associated with the learned word pairs were delivered 3 times each, for a total number of 54 auditory stimulations. The duration of each sound was 6 seconds, and the inter-stimulus interval randomly ranged 6 to 12 seconds. In the TMR-anodal left tDCS condition, participants (N = 17) were administered the TMR protocol and received in parallel tDCS with the anode over the left DLPFC and the cathode over the right DLPFC. TMR-anodal right tDCS condition (N = 20) was identical to TMR-anodal left tDCS, except that the polarity of the electrodes was reversed (right anode/left cathode on tDCS). In the TMR-sham tDCS condition (N = 16), tDCS was delivered only for 15 seconds at the beginning of the TMR procedure.

**Figure 1.**
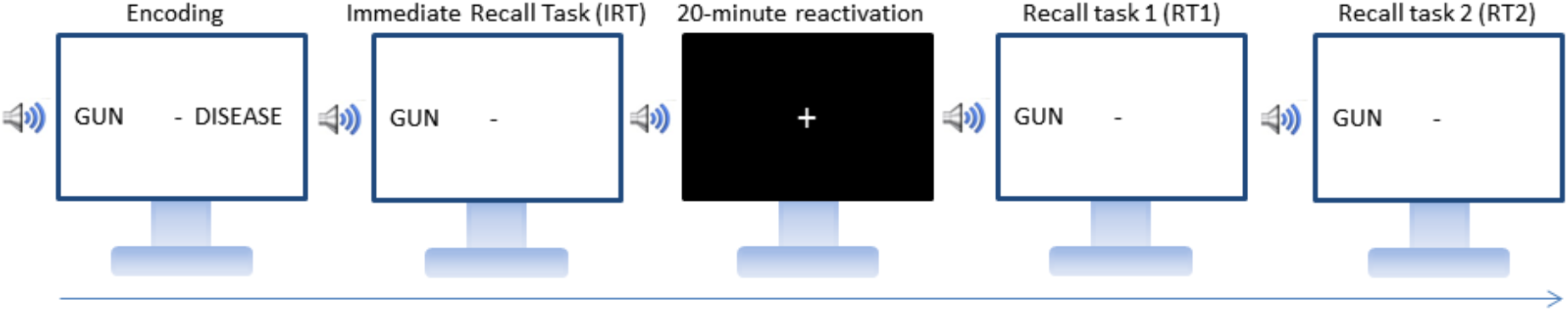
Experimental design. Participants learned word pairs associated with emotionally compatible sounds until reaching 75% correct recall in the immediate cued recall task (IRT). During the 20-minute reactivation period in a wakeful resting state, the sounds associated with half of the neutral and negative word pairs were replayed three times each alone (TMR-only) or concurrently with excitatory tDCS on the left (TMR-anodal left tDCS) or right (TMR-anodal right tDCS) hemisphere or in a sham tDCS condition (TMR-sham). Memory for all learned word pairs was then tested in a cued recall task immediately after (RT1) and one week later (RT2).

Immediately after the 20-minutes stimulation period, participants were administered a cued recall task (RT1) on all word pairs presented in the learning session. RT1 was identical to the immediate recall task (IRT), except that participants did not receive any feedback on the correctness of their responses and there was no cut-off score. One week later, participants were administered a second cued recall session (RT2) in identical conditions and at the same time of the day than RT1.

Subjective sleepiness and objective vigilance were assessed at the beginning of the learning (KSS 1, PVT 1), RT1 (KSS 2, PVT 2) and RT2 (KSS 3, PVT 3) sessions using the Karolinska Sleepiness Scale (KSS, Akerstedt & Gillberg, 1990) and the 10-min version of the Psychomotor Vigilance Task (PVT, Dinges & Powell, 1985) respectively.

### 2.4. Transcranial Direct Current Stimulation

A pair of saline-soaked sponge electrodes (50 × 70 mm) was positioned on the scalp at F3 and F4 locations (Figure 2) according to the 10-20 international system for electrode placement, determined using the Beam F3 location system (Beam, Borckardt, Reeves, & George, 2009). Direct current stimulation was delivered using a DC-Stimulator Plus (Rogue Resolutions, Cardiff, UK) operated in accordance with published safety guidelines (Antal et al., 2017). In the left tDCS condition, stimulation was anodal (excitatory) on F3 and cathodal (inhibitory) on F4. In the right tDCS condition, polarity was reversed (anode at F3 and cathode at F4). Current intensity was set at 1 milliamp, corresponding to a current density of 0.029 mA/cm^2^. The stimulation ascended during a 10 seconds ramp up to 1 mA, then stabilized within a 20-minute plateau and faded out to 0 mA within 10 seconds. The plateau lasted only 15 seconds in the sham tDCS condition, then stimulation was discontinued after fading out. Participants were blind with respect to the type of stimulation.

**Figure 2.**
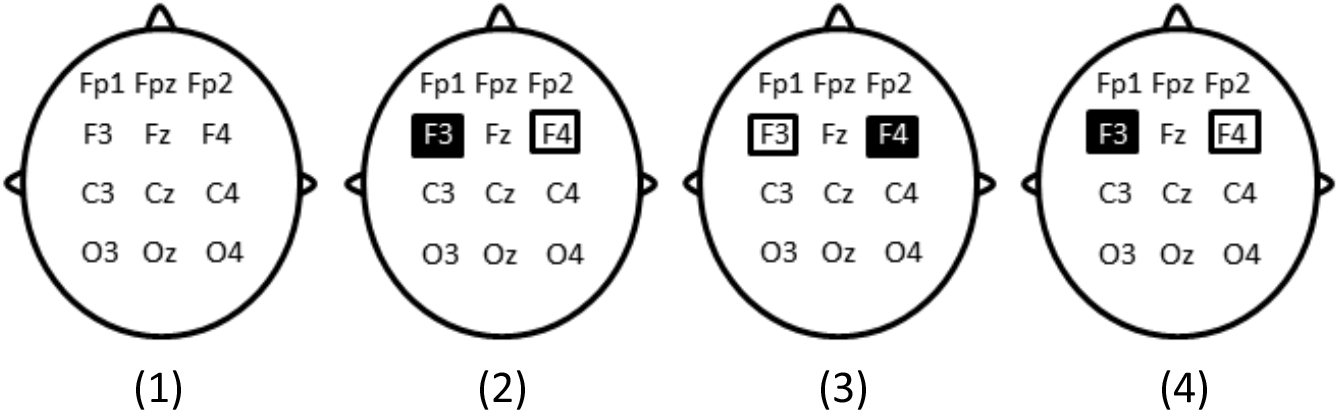
Schematic representation of electrodes’ position in the four conditions: (1) TMR-only, no electrode positioned. (2) TMR-anodal left tDCS, anodal stimulation on the left DLPFC and cathodal stimulation on the right DLPFC for 20 minutes. (3) TMR-anodal right tDCS, cathodal stimulation of the left DLPFC and anodal stimulation of the right DLPFC for 20 minutes. (4) Sham condition, anodal stimulation of the left DLPFC and cathodal stimulation of the right DLPFC for 15 seconds. Dark rectangle depicts the anode and light rectangle depicts the cathode.

## 3. Results

### 3.1. Sleepiness and vigilance

A repeated measure ANOVA conducted on subjective sleepiness (KSS) scores with within-subjects factor Moment (KSS 1 (learning) vs. KSS 2 (RT1) vs. KSS 3 (RT2)) and between-subjects factor Condition (TMR-anodal left tDCS, TMR-anodal right tDCS, TMR-sham tDCS and TMR-only) failed to disclose significant differences in sleepiness scores from learning (3.42 ± 1.7) to retrieval (RT1 = 3.19 ± 1.1, RT 2 = 3.45 ± 1.6, *F*(3, 65) = 0.48, *p* = .627). The main effect of Condition and the Condition by Moment interaction were also non-significant (all *ps* > .370).

Similar analyses were conducted on two PVT parameters, i.e. the coefficient of variation and the reciprocal RT (mean 1/RT; Basner and Dinges, 2011). Again, no differences were evidenced between the encoding and the recall session either using the coefficient of variation (PVT 1 = 0.168 ± 0.08 vs. PVT 2 = 0.159 ± 0.06, PVT 3 = 0.162 ± 0.05 *F*(3, 65) = 0.81, *p* = .396), or the reciprocal RT (PVT 1 = 0.0029 ± 0.0004 vs. PVT 2 = 0.0030 ± 0.0005, PVT 3 = 0.0030 ± 0.0003, *F*(3, 65) = 0.49, *p* = .488). The main effect of Condition and the Condition by Moment interaction were also non-significant (all *ps* > .311).

Altogether, these results indicate no significant differences between the encoding and recall sessions in sleepiness and vigilance states, and that these states were not modulated by the intermediate administration of tDCS and TMR.

### 3.2. Pre-stimulation Learning Session (IRT)

A repeated measure ANOVA computed on the number of correctly retrieved word pairs at the end of learning (IRT) task with within-subject factors Cueing (Cued vs. Non-Cued) and Emotion (Neutral vs. Negative) and between-subject factor Condition (TMR-anodal left tDCS, TMR-anodal right tDCS, TMR-sham tDCS and TMR-only) only disclosed a main effect of Emotion (*F*(1, 65) = 35.16, *p* < .001) with better learning for neutral (mean ± standard error, 89.81 ± 0.98%) than negative word pairs (79.65 ± 1.1%; Figure 3), although performance was above the 75 % cut-off score for both categories. No significant differences in retrieval score were observed between the word pairs to be subsequently cued and the word pairs not to be subsequently cued (*F*(1, 65) = 0.13, *p* = .717), nor between the four conditions (*F*(3, 65) = 0.93, *p* = .433). All other main and interaction effects were non-significant (all *ps* > .174). Hence, pre-stimulation conditions were similar across the four experimental conditions.

**Figure 3.**
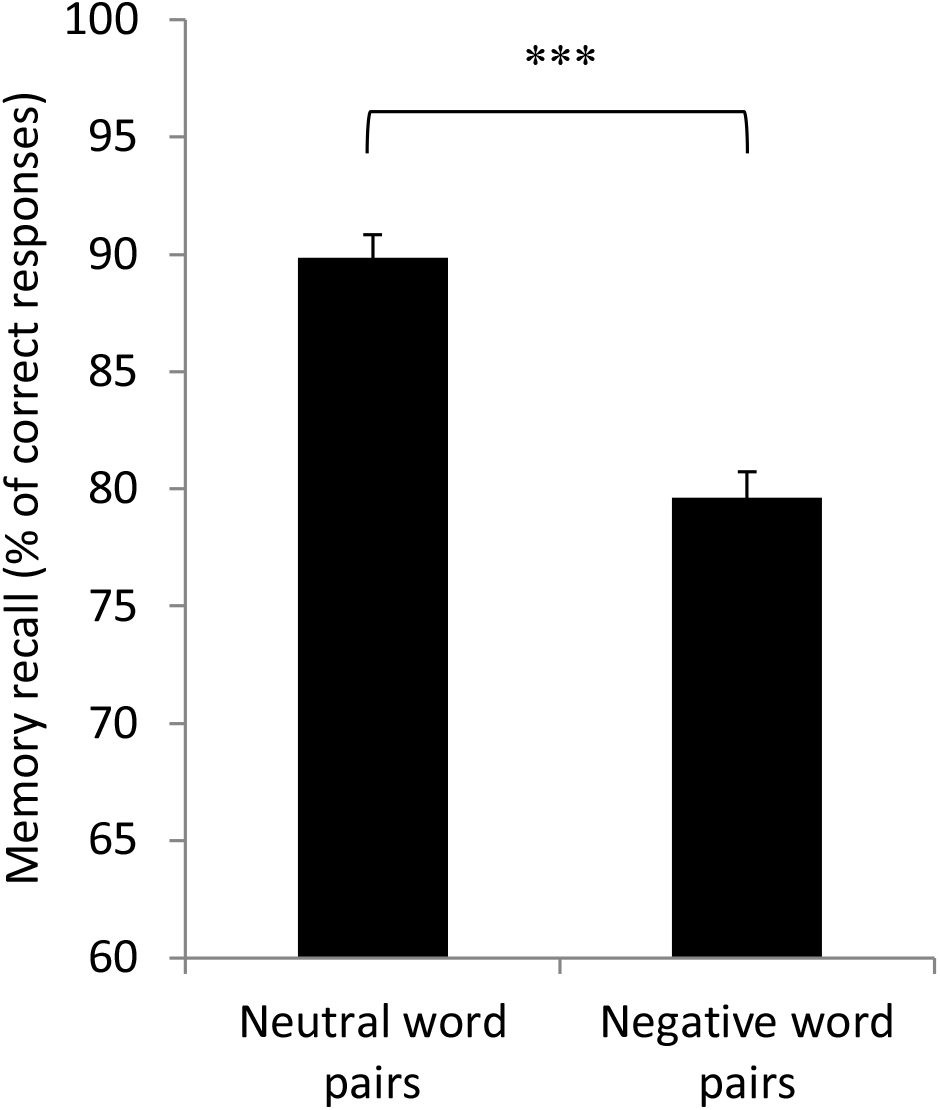
Memory performance for neutral and negative word pairs at the immediate recall task (IRT), expressed in percentage of correct responses. Error bars illustrate standard error. *** *P* < .001.

### 3.3. Immediate post-stimulation testing session (RT1)

A repeated measure ANOVA computed on the percentage of correctly recalled word pairs with within-subject factors Emotion (Neutral vs. Negative Word pairs) and Cueing (Cued vs. Non-Cued) and between-subject factor Condition (TMR-anodal left tDCS, TMR-anodal right tDCS, TMR-sham tDCS and TMR-only) disclosed a main effect of Condition (*F*(3, 65) = 4.58, *p* = .006). There was a main effect of Cueing (*F*(1, 65) = 6.78, *p* = .011) with higher recall for Cued 82.90 ± 1.7%) than Non-Cued (79.34 ± 1.7%) word pairs. In addition, there was a significant Cueing by Condition interaction (*F*(3, 65) = 4.41, *p* = .006; Figure 4). The Cueing by Emotion interaction effect was not significant (*F*(3, 65) = 0.69, *p* = .410) nor the Condition by Cueing by Emotion interaction effect (*F*(3, 65) = 0.64, *p* = .595). Similarly, the Condition by emotion was not significant (*F*(1, 65) = 0.86, *p* = .467).

**Figure 4.**
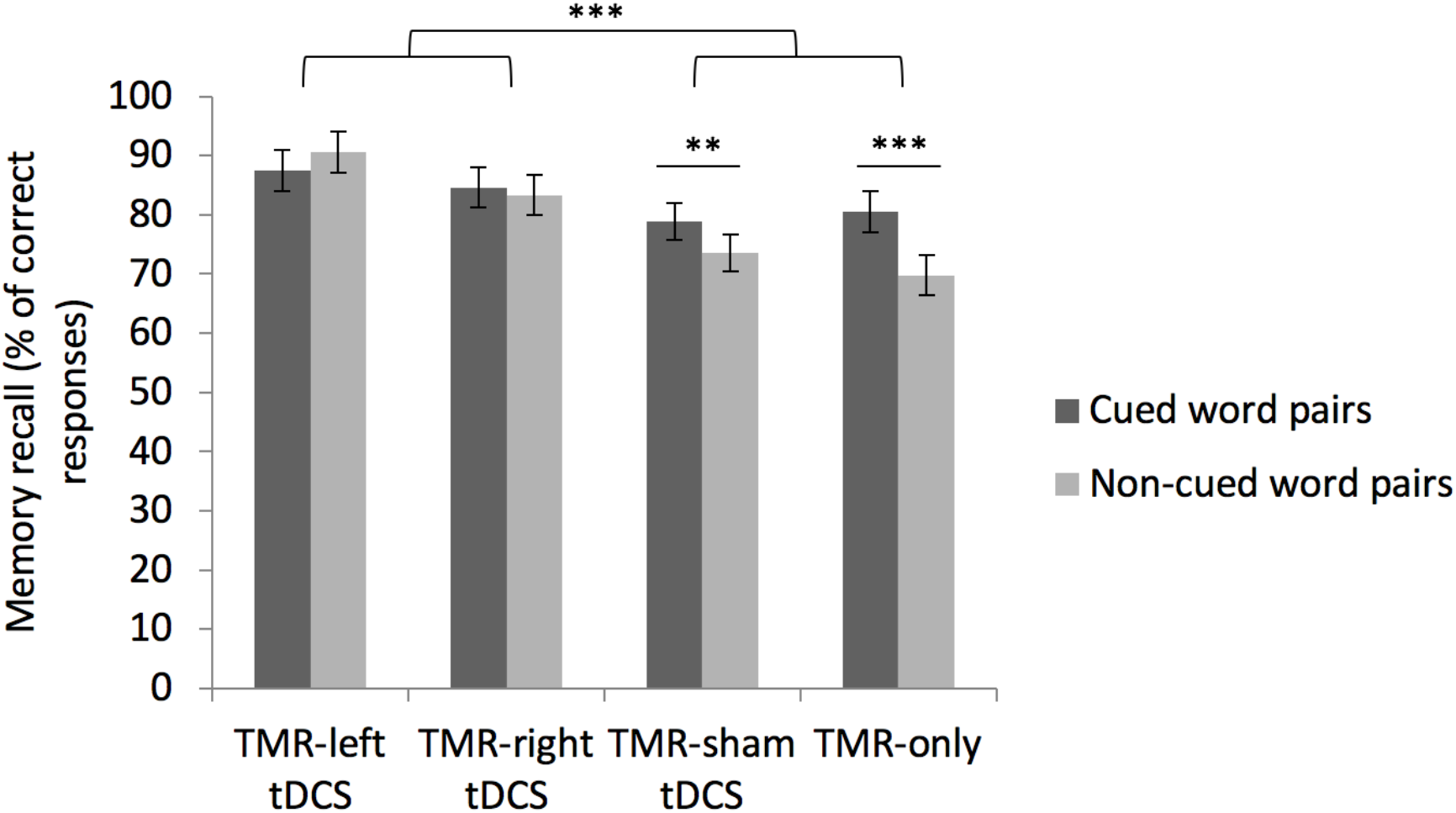
Memory performance for cued and non-cued word pairs at the first Recall task (RT1) in the four conditions. Error bars illustrate standard error. \*\**p* < .01, \*\*\**p* < .001.

Planned comparisons conducted on the significant between-subject factor Condition showed that memory decline was significantly lower in conditions in which participants received real electrical stimulation (TMR-anodal left tDCS and TMR-anodal right tDCS) than in conditions in which no or a sham electrical stimulation was applied (TMR-sham tDCS and TMR-only; F(1, 65) = 12.58, *p* < .001). A separate comparison between tDCS real (TMR-anodal left tDCS and TMR-anodal right tDCS) and sham (TMR-sham tDCS) conditions confirmed better performance in the tDCS real conditions (*F*(1, 65) = 8.26, *p* = .005). The comparison between TMR–anodal left tDCS and TMR-anodal right tDCS conditions was non-significant (*F*(1, 65) = 1.33, *p* = .252). Likewise, no significant differences were evidenced between the TMR-sham tDCS and the TMR-only conditions (*F*(1, 65) = 0.06, *p* = .800).

Additional planned comparisons including the significant within-subject factor Cueing disclosed a significant Cueing effect in conditions in which no or a sham electrical stimulation was applied (TMR-sham tDCS and TMR-only; *F*(1, 65) = 17.916, *p* < .001), but not in conditions in which participants received real electrical stimulation (TMR-anodal left tDCS and TMR-anodal right tDCS; *F*(1, 65) = 0.22, *p* = .646). A direct comparison between TMR-sham tDCS and TMR-only conditions indicates that the Cueing effect was not significantly different (*F*(1, 65) = 2.09, *p* = .153).

Altogether, these results suggest that tDCS significantly benefitted memory consolidation irrespective of the side of stimulation and of TMR effects. A benefit of targeted memory reactivation was only observed in the TMR-sham tDCS and TMR-only conditions. Finally, the emotional valence of the word pairs did not elicit any main or interaction effects, suggesting that neutral and negative memories equally benefitted from TMR, and that tDCS laterality did not interact with the emotional valence of the material.

### 3.4. Long term memory consolidation (RT2)

Similarly, memory performance at delayed recall (one week later) was computed on the number of correctly retrieved learned word pairs, expressed in percentage. The ANOVA with within-subject factors Emotion (Neutral vs. Negative Word pairs) and Cueing (Cued vs. Non-Cued) and between-subject factor Condition (TMR-anodal left tDCS, TMR-anodal right tDCS, TMR-sham tDCS and TMR-only) disclosed a main effect of emotion (*F*(1, 64) = 32.44, p < .001). All other main and interaction effects were non-significant (all *ps* > .185; Figure 5). Average forgetting across conditions was also significantly more pronounced in RT2 than RT1 (*F*(1, 64) = 234.31, *p* < .001).

**Figure 5.**
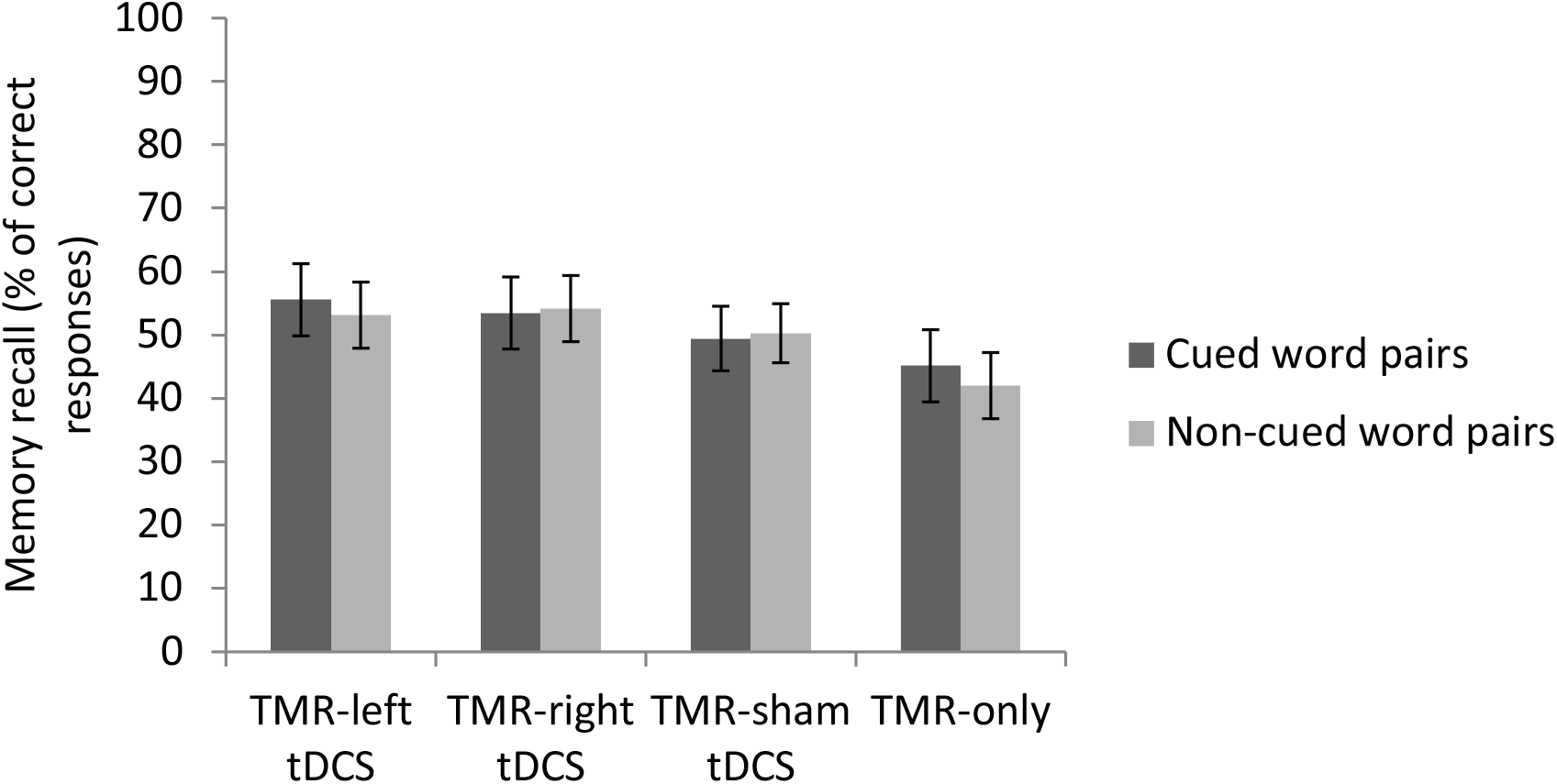
Memory forgetting for cued and non-cued word pairs from Immediate (IRT) to delayed (RT2) recall in the four conditions. Error bars illustrate standard error.

## 4. Discussion

In the present study, we first tested the hypothesis that providing auditory reminders (i.e. a TMR procedure) during a wakeful rest period would enhance memory consolidation for targeted items. Our results confirm a significant benefit of auditory reminders, but only in the TMR-only and TMR-sham tDCS conditions; that is in the absence of effective electrical stimulation. Secondly, we hypothesized that electrical stimulation of the DLPFC would improve the retention of word pairs and reinforce TMR effects. Our results partially fulfilled these predictions. Indeed, retrieval was significantly better in the TMR-anodal left and TMR-anodal right tDCS conditions than in the no- and sham stimulation conditions, confirming a beneficial effect of tDCS on memory consolidation, but this effect was generalized to all learned items irrespective of the TMR procedure, i.e. against the hypothesis that tDCS potentiates the selective enhancing benefits of TMR on memory. Finally, we tested the hypothesis that anodal excitatory stimulation of the right DLPFC associated with cathodal inhibitory stimulation of the right DLPFC would enhance the benefits of TMR for negative word pairs, eventually leading to greater benefits for negative than neutral cued word pairs. Results did not evidence polarity-dependent hemispheric effects on the consolidation of negative memories.

### 4.1. The benefits of auditory cueing on memory consolidation

As stated above, there was a selective memory enhancement for cued as compared to non-cued word pairs in the TMR-only tDCS and TMR-sham tDCS conditions, while this effect was completely abolished in both TMR-anodal left tDCS and TMR-anodal right tDCS conditions. These results partially corroborate our primary hypothesis of a benefit of TMR for the consolidation of targeted items in memory. However, the fact that the cueing advantage was totally absent in both TMR-anodal left and TMR-anodal right tDCS conditions suggests that the selective effect of TMR was actually overshadowed by the global effect of tDCS. Indeed, no forgetting was observed in the recall task immediately after stimulation (RT1) for both cued and uncued items in both tDCS conditions, whereas forgetting was actually more pronounced for uncued than cued items in the TMR-only and TMR-sham tDCS conditions. Hence, the cueing benefit may have been abolished in effective tDCS conditions, possibly due to a better global memory retention masking the benefits of TMR.

It was shown already that different factors might determine the effectiveness of TMR. For instance, auditory cueing at wake was found mostly beneficial to rescue low reward value spatial stimuli from forgetting, while it did not impact on high reward value stimuli (Oudiette et al., 2013). In this latter study, low reward value stimuli actually also exhibited lower learning accuracy than high reward value items before the TMR intervention, suggesting that TMR at wake is mostly efficacious when learning levels are moderate. Similarly, a benefit of auditory cueing during sleep on memory for object locations was found only for items that were not already highly accurate before the intervention (Creery, Oudiette, Antony, & Paller, 2015). Hence, initially high encoding levels might explain why other studies failed to evidence even a moderate benefit of TMR at wake. Notwithstanding, negative items were less efficiently learned than neutral ones in our present study, and still did not benefit more from TMR or tDCS, as there was no main or interaction effect of the emotional valence of the learned stimuli. Since both negative and neutral items were already learned above 75% accuracy in our study, it might be that this initial difference was insufficient to trigger differential effects either of TMR or tDCS.

### 4.2. The effects of tDCS on memory consolidation

As expected, electrical stimulation of the DLPFC (TMR-anodal left and TMR-anodal right tDCS) led to significantly better recall performance as compared to the TMR-only and TMR-sham tDCS conditions. Therefore, 20 minutes of direct current stimulation over the left or right DLPFC during resting wakefulness can boost memory consolidation for declarative verbal material. These results are fitting with the report that tDCS over the left DLPFC at wake strengthens episodic memories and reduces further forgetting (Sandrini et al., 2014; Sandrini et al., 2019). Inconsistently however, Kirov and colleagues (Kirov, Weiss, Siebner, Born, & Marshall, 2009) found that transcranial slow (0.75 Hz) oscillation stimulation (tSOS) applied 20 minutes after learning did not improve the retention of declarative memories, although it increased endogenous EEG slow oscillation as well as theta activity. However, methodological differences (electrode size and montage) and study design (postlearning electrical stimulation delayed by 20 minutes) might explain these discrepancies. In our study, the reactivation of memories triggered by the TMR procedure might have contributed to the memory benefits following tDCS. Because the TMR procedure reinstated the activation of the mnemonic traces, the electrical brain stimulation might have stabilized memory through a process of consolidation or reconsolidation. Accordingly, Javadi and Cheng (2013) found that anodal stimulation of the left DLPFC during a consolidation interval improved memory performance, but only when the memory traces were reactivated during the stimulation using an old-new recognition task. Together, these results point out the combination of memory reactivation procedures and electrical brain stimulation as a promising method to enhance memory consolidation.

The exact mechanisms by which tDCS improves memory consolidation when applied during a resting state following learning are still unclear, even if we have increasing knowledge about the effects of tDCS on cortical excitability (Nitsche et al., 2003; Yavari, Jamil, Mosayebi Samani, Vidor, & Nitsche, 2018). Neurophysiological studies showed that anodal tDCS induces cortical excitability and enhances NMDA receptors plasticity (Liebetanz, Nitsche, Tergau, & Paulus, 2002; Nitsche et al., 2003; Nitsche & Paulus, 2001; Stagg, Antal, & Nitsche, 2018). Memory formation is also known to depend on changes in synaptic strength such as synaptic tagging and long term potentiation (Wang, Redondo, & Morris, 2010). Thus, the benefits of tDCS over the DLPFC might primarily stem from a direct modulation of synaptic plasticity processes within prefrontal areas, in itself favouring memory reorganization, eventually leading to improved recall accuracy. Secondly, as mentioned above, memory consolidation processes are thought to rely on the offline reactivation of learning-related neural activity (Peigneux et al., 2006; Tambini & Davachi, 2019; Wamsley, 2019). The reactivation of declarative, hippocampus-dependent memories through a dialogue between hippocampal and neocortical areas is possibly mediated by slow oscillatory activity at wake (Brokaw et al., 2016) like during sleep (Buzsaki, 1996). Transcranial DCS-related increased excitability in prefrontal regions could also favour connections with remote memory-related areas. For instance, anodal tDCS over the primary motor cortex was shown efficient to increase functional coupling with subcortical thalamus regions (Polania, Paulus, & Nitsche, 2012). In the present case, it can be speculated that increasing brain excitability within prefrontal areas might boost connections with hippocampal areas, thereby increasing the susceptibility of memory traces to be reactivated. However, we cannot conclude for a specific involvement of the DLPFCs as our study did not include a control condition with the stimulation of a region not supposed to be involved in these processes, such as the primary motor cortex.

Hemispheric specialisation of the DLPFC in learning and memory is a matter of debate in literature. It was proposed that the left DLPFC mostly supports encoding while the right DLPFC mostly supports retrieval (Tulving, Kapur, Craik, Moscovitch, & Houle, 1994), alternatively that the left DLPFC supports the consolidation of verbal material while the right DLPFC processes non-verbal material (Cabeza & Nyberg, 2000). Although our results support the idea that tDCS over the DLPFC during a consolidation episode benefits memory for verbal declarative material, we did not find an effect of tDCS polarity on behavioural outcomes. As well, prior studies led to contrasting results regarding an asymmetric involvement of the DLPFC. As stated above, Sandrini et al. (2014) showed that 15 minutes of tDCS over the left DLPFC improves recall for verbal material. Several studies also support the proposal of a specialisation of the left DLPFC in memory consolidation (Javadi & Cheng, 2013; Javadi & Walsh, 2012; Sandrini et al., 2014; Westphal et al., 2019). Nonetheless, an implication of the right DLPFC in the reactivation and consolidation of episodic memories was evidenced in a fMRI study (Diekelmann et al., 2011) with increased activity in the right lateral prefrontal cortex after reexposure to an odour associated with the context of learning. Likewise, 1 Hz repetitive transcranial magnetic stimulation (rTMS) applied after memory reactivation over the same region was found to improve retention of episodic memories (Sandrini, Censor, Mishoe, & Cohen, 2013).

### 4.3. Consolidation of emotional memories and lateralisation of tDCS polarity

We hypothesized that anodal excitatory stimulation of the right DLPFC (with cathodal inhibitory stimulation of the left DLPFC) would increase the selective enhancement effect of TMR for negative word pairs. However, our results did not confirm a polarity-dependent effect of tDCS for the consolidation of negative items. Therefore, we cannot conclude that the polarity of tDCS applied during a consolidation interval modulates TMR for negative memories. Like for a global TMR effect however, it is possible that the main and powerful effect of tDCS on overall memory performance masked a specific effect of tDCS polarity on the consolidation of negative word pairs.

Although this possibility should be investigated in further studies, others similarly failed to evidence inter-hemispheric dissociations between left and right DLPFC in the processing of negative emotions, or even obtained unexpected opposite effects. For instance, anodal tDCS over the left DLPFC was found to facilitate the recognition of negative and positive facial expressions (with a more pronounced benefit for positive emotions), while it did not impact the emotional state of the participants (Nitsche et al., 2012). Similarly, Penolazzi et al. (2010) found that right anodal/left cathodal stimulation over fronto-temporal regions facilitated recall for pleasant images, whereas left anodal/right cathodal stimulation improved recall for unpleasant images. Notwithstanding, our results should be taken cautiously as neutral word pairs were better encoded than negative ones at the end of learning already. Superior learning for neutral items is surprising given that arousing negative emotional material is usually better encoded (Kilpatrick & Cahill, 2003). Indeed, most studies having investigated the influence of emotional valence in the time course of memory consolidation found emotional memories to be usually better remembered over time (Ochsner, 2000), probably because the amygdala mediates the organization of memories in the hippocampus and the neocortex (de Voogd, Fernandez, & Hermans, 2016; Liu et al., 2016). In our current study, it cannot be excluded that the arousing effect of the negative sounds associated with the word pairs was too high, which would have shifted the participants’ attention toward the sounds, to the detriment of the associated negative word pairs. Prior studies also found a specific deleterious effect of arousing negative emotions on associative learning (Bisby & Burgess, 2013; Guez, Saar-Ashkenazy, Mualem, Efrati, & Keha, 2015; Nashiro & Mather, 2011). Such attentional shift might have interfered with encoding and later retrieval processes, possibly masking a specific effect of right anodal/left cathodal stimulation of the DLPFC.

### 4.4. No long-term benefits of TMR and tDCS

A 30 to 40 % forgetting was similarly observed in all conditions when retested one week later, suggesting that wake TMR and tDCS-related benefits on memory consolidation are short lived, contrarily to prior reports (e.g. Floel et al., 2012). It is possible that the high level of memory performance achieved at the first recall session may have masked the long-term benefit of TMR and tDCS in our study. In the Flöel et al. study, elderly subject might have had more room for memory improvement, increasing the potential tDCS-related memory improvement on the long-term. Another possible explanation is that presenting auditory reminders during a period of wakeful rest initially boosts the associated memories but concomitantly puts those into a more labile condition more susceptible to external interference eventually leading to forgetting. For instance, re-exposure to a contextual odour at wake was shown to impair the retrieval of images location (Diekelmann et al., 2011), probably because memories are more labile after reactivation and need to be reconsolidated (Javadi & Cheng, 2013; Nader, 2003; but see Crossman, Bartl, Soerum, & Sandrini, 2019). Additionally, the time interval between learning and delayed recall was seven days, without any reminders. It is possible that cueing memories over consecutive days might have led to identifiable benefits on the long-term. Further studies are needed to disentangle the temporal effects of both tDCS and TMR techniques.

## 5. Conclusions

In summary, we have shown in the present study that TMR during a wakeful resting period benefits short-term memory consolidation. However, concomitant tDCS either on right or left DLPFC gave rise to much higher but unspecific memory enhancement, hence abolishing or at least overshadowing the TMR advantage. Finally, our results did not evidence a polarity-dependent hemispheric effect of tDCS on the consolidation of emotional negative memories. Noticeably, stimulation of the DLPFC during a 20-minute period following learning was found beneficial for the consolidation of verbal declarative memories. By increasing cortical excitability in prefrontal areas, tDCS might favour the hippocampo-cortical dialogue subtending memory consolidation processes.

## Acknowledgments

The study was supported by the FRS-FNRS Projet de recherche (PDR) T.0109.13 and FNRS-FWO Excellence of Science (EOS) MEMODYN project. At the time of the study, Médhi Gilson was FRS-FNRS Research Fellow.

## Author Contributions

Médhi Gilson was the leading researcher for this study. Michael Nitsche contributed to the experimental design and to the redaction of the article. Philippe Peigneux contributed to the experimental design, to data analysis and to the redaction of the article.

## Conflicts of Interest

The authors declare no conflict of interest.

## Notes

### Competing Interest Statement

The authors have declared no competing interest.

